# A long-distance inhibitory system regulates haustoria numbers in parasitic plants

**DOI:** 10.1101/2024.12.19.629485

**Authors:** Anna Kokla, Martina Leso, Jan Simura, Cecilia Wärdig, Marina Hayashi, Naoshi Nishii, Yuichiro Tsuchiya, Karin Ljung, Charles W. Melnyk

**Author notes:** Corresponding author: Charles W. Melnyk, Department of Plant Biology, Linnean Center for Plant Biology, Swedish University of Agricultural Sciences, Almas allé 5, 756 51, Uppsala, Sweden. These authors contributed equally to this work (Anna Kokla, Martina Leso).

## Abstract

The ability of parasitic plants to withdraw nutrients from their hosts depends on the formation of an infective structure known as the haustorium. How parasites regulate their haustoria numbers is poorly understood, and here, we uncovered that existing haustoria in the facultative parasitic plants *Phtheirospermum japonicum* and *Parentucellia viscosa* suppressed the formation of new haustoria on distant roots. Using *Phtheirospermum japonicum,* we found that this effect depended on the formation of mature haustoria and could be induced through the application of external nutrients. To understand the molecular basis of this root plasticity, we analyzed hormone response and found that existing infections upregulated cytokinin responsive genes first at the haustoria and then more distantly in *Phtheirospermum* shoots. We observed that infections increased endogenous cytokinin levels in *Phtheirospermum* roots and shoots, and this increase appeared relevant since local treatments with exogenous cytokinins blocked the formation of both locally and distantly formed haustoria. In addition, local overexpression of a cytokinin degrading enzyme in *Phtheirospermum* prevented this systemic inter-haustoria repression and increased haustoria numbers locally. We propose that a long-distance signal produced by haustoria negatively regulates future haustoria, and in *Phtheirospermum*, such a signaling system is mediated by a local increase in cytokinin to regulate haustoria numbers and balance nutrient acquisition.

## Introduction

Plant parasitism has evolved at least 12 independent times resulting in thousands of different parasitic plant species that share a common feature: an infective structure known as the haustorium (1). Several parasitic plant species are important agricultural pests including *Striga* and *Cuscuta* that cause major economic losses every year (2, 3). These two species are obligate parasitic plants that depend entirely on their hosts for survival but many other species are facultative parasites that are independent of their hosts and instead parasitize under the right conditions (4). Many parasitic plants have important ecological roles, for example they contribute to the spread of other species and biodiversity by parasitizing dominant species within their ecosystem (5). Using the haustorium, parasitic plants invade their hosts and form xylem connections, and some species also form phloem connections, to uptake water, nutrients, hormones and RNAs (6–9).

Among the most studied parasitic plants is the facultative root parasite *Phtheirospermum japonicum*. *Phtheirospermum* initiates pre-haustoria formation in response to haustorium inducing factors (HIFs) released by hosts such as 2,6-dimethoxy-1,4-benzoquinone (DMBQ) (10–12). Following initiation, the pre-haustorium attaches to the host root with the help of specialized root hairs (13), penetrates the host tissues using cell wall-modifying enzymes (14) and matures to form a haustorium with xylem connections known as the xylem bridge (15). In addition to HIFs, endogenously produced compounds such as auxin and ethylene are important for successful haustoria formation and xylem bridge development (15, 16). Negative regulators of haustoria formation are less well known but include exogenous nitrogen which, in *Phtheirospermum*, suppresses haustoria numbers via the upregulation of abscisic acid (17).

Plant hormones play important roles in the regulation and development of haustoria. One such hormone, cytokinin, is synthesized by ISOPENTENYLTRANSFERASE (IPT) and LONELY GUY (LOG) proteins and these compounds can either act locally or move long distances where they are perceived by ARABIDOPSIS HISTIDINE KINASE (AHK) receptors to activate type B ARABIDOPSIS RESPONSE REGULATOR (ARR) transcription factors and cytokinin degrading CYTOKININ OXIDASE (CKX) enzymes (18). Cytokinin levels increase at the site of *Phtheirospermum* infection and move from the parasite to the host to induce host root expansion (7, 19). Cytokinins also play important roles in other symbiotic relationships such as during parasitism by nematodes, when nematodes release cytokinins to activate cell division and form syncytium feeding sites (20). Similarly, cytokinins are essential for the formation of symbiotic structures called nodules that form between legumes and nitrogen fixing bacteria. Such symbiotic relationships are often characterized by both positive and negative regulators that help balance the numbers of symbiotic structures to optimize resources use by the plant, a process which in legumes is known as auto-regulation of nodulation (AON) (21).

Although the regulation of symbiosis is well known in legumes, the endogenous signals that regulate parasitic plant infection remain poorly characterized and it is unknown whether the numbers of haustoria that form are regulated by the parasite. Here, we identified a system in the Orobanchaceae family members *Parentucellia viscosa* and *Phtheirospermum japonicum* whereby existing haustoria control the formation of new haustoria via a systemic repressive signal. We investigated the increase in cytokinins by *Phtheirospermum* during infection and found that they serve an inhibitory role for the formation of new haustoria found on both local and distant roots. We propose that cytokinin regulates haustoria formation as part of a long-distance repressive signal, thus allowing the parasite to control haustoria numbers and infection plasticity.

## Results

### A systemic signal controls haustoria numbers

Plants involved in nutrient-acquiring symbioses often regulate the extent of symbiosis (21, 22) but whether such regulation exists in plant parasitism is unknown. To investigate whether plant parasites control their number of nutrient-acquiring organs, the haustoria, we infected *Arabidopsis* with *Phtheirospermum* at day 0, followed by a second infection with a new host added on the same *Phtheirospermum* root 10 days later (Fig. 1A). This second infection showed significantly fewer haustoria and reduced xylem bridge development compared to the first infection. To exclude age or starvation time effects, we performed first infections on plants at 10 days. These infections showed intermediate haustoria numbers but no difference in xylem bridge development compared to first infections at 0 days post infection (dpi). To investigate whether such a phenomenon might operate over long distances, we developed a “split-root” system where *Phtheirospermum* roots were separated in two sides (Fig. 1B). When the host was added at the same time on both sides (Day 0-Day 0), both sides of the *Phtheirospermum* root system formed the same number of haustoria (Fig. 1C). We then added the hosts to one side at day 0, and waited 3, 5, 7 or 10 days before adding the hosts to the second side. While the Day 0 side formed the same number of haustoria as the Day 0-Day 0 control, the second side showed a progressive reduction in haustoria numbers (Fig. 1C). The age and starvation time control (Day 10-Day 10) had comparable numbers of haustoria to the Day 0-Day 0 plants (Fig. 1C). The haustorium inducing factor DMBQ causes pre-haustoria to form in *Phtheirospermum* that do not mature to form xylem bridges (13), so we pre-treated one side with DMBQ and found it did not inhibit the formation of haustoria on the distant side (Fig. 1D). In addition, removing the host added on the first side after 5 days of infection inhibited haustoria formation on the second side (Supplemental Fig. S1A). Together, these data suggested that mature haustoria were needed for systemic inhibition and that the inhibitory signal lasted for several days even if hosts were removed. Nutrients can regulate symbiosis, and in *Phtheirospermum*, exogenous nitrogen inhibits haustoria formation via ABA signaling (17). Applying 10.3 mM NH_4_NO_3_ to one side of the split-root setup inhibited haustoria both locally and systemically suggesting that nitrogen could function as part of a negative regulatory signal (Fig. 1E). However, ABA treatment inhibited haustoria formation only locally indicating ABA was not mobile or part of the systemic signaling process (Supplemental Fig. S1B), thus leaving the identity of the endogenous signal unresolved. To test whether systemic inhibition was found in other plant parasites, we performed split plates assays with *Parentucellia viscosa* and found that, similar to *Phtheirospermum*, existing haustoria suppressed the formation of new haustoria (Fig. 1F) indicating this regulatory system was not unique to *Phtheirospermum*.

**Figure 1:**
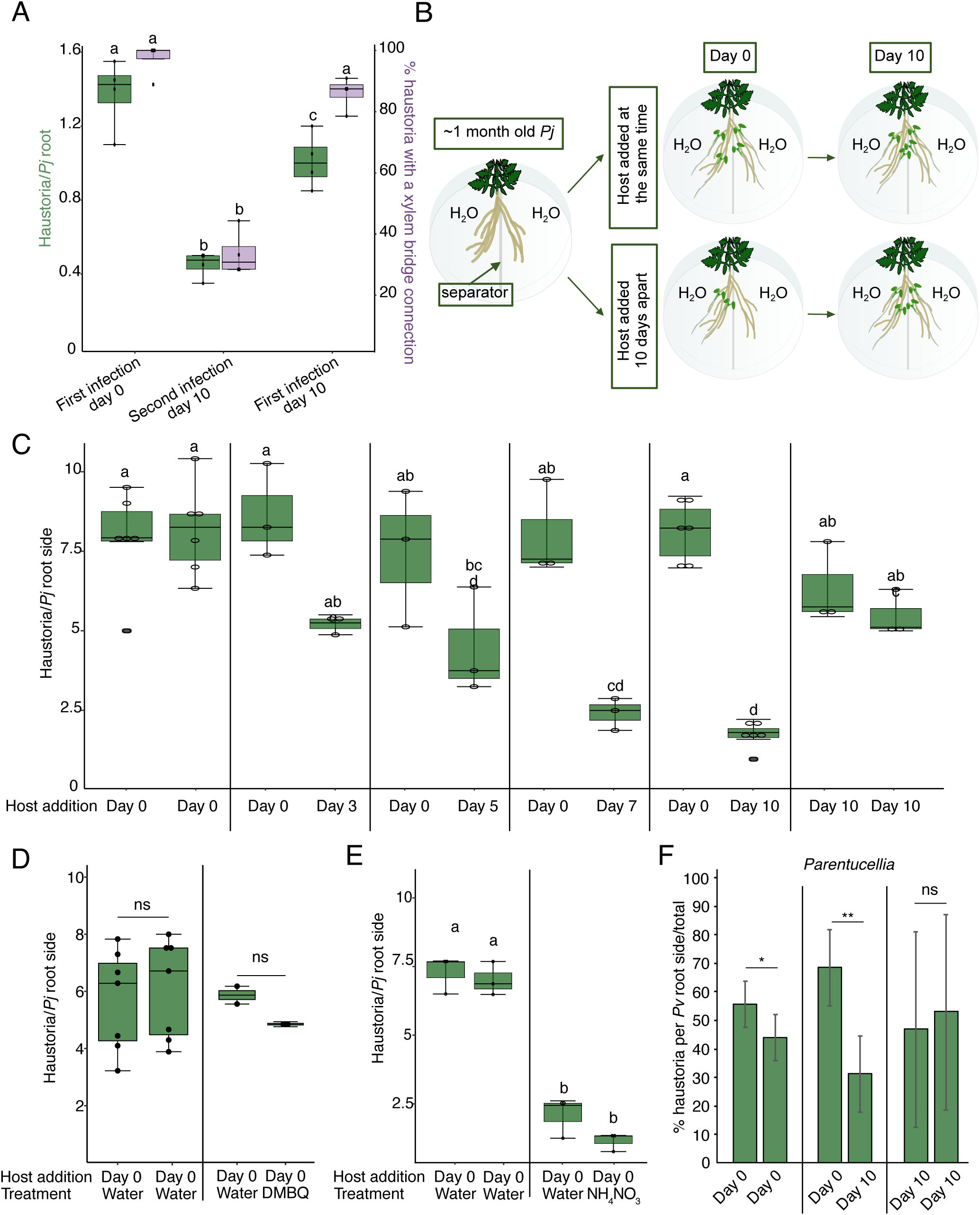
Haustoria numbers are controlled by systemic signaling. **A)** Numbers of haustoria and percentage of haustoria with xylem bridges per *Phtheirospermum* with infection at day 0 followed by a second infection at day 10 on the same root. Control is first infection at day 10. The haustoria were evaluated 7 days after each infection. (n = 3 replicates) **B)** Drawing of the split-root experimental setup. **C)** Average number of haustoria per *Phtheirospermum* root side in a split-root setup on water agar, with host added on one side at 0 days post infection (Day 0) and on the other side 3, 5, 7 or 10 days later, or on both sides at 0 or 10 days post infection. (n = 3-6 replicates) **D)** Average number of haustoria per *Phtheirospermum* root side in a split-root setup on water agar or 10 μM DMBQ, with host added on both sides at day 0. (n = 2-6 replicates) **E)** Average number of haustoria per *Phtheirospermum* root side in a split-root setup on water agar or 10.3 mM NH_4_NO_3_, with host added on both sides at day 0. (n = 3 replicates) **F)** Percentage of haustoria per *Parentucellia* root side in a split-root setup on water agar, with host *Arabidopsis* added on one side at 0 days post infection (Day 0) and on the other side 10 days later, or on both sides at 0 or 10 days post infection. (n = 3-4 plants, two-tailed Student’s t-test, * p < 0.05, ** p < 0.01, ns= not significant) **A,C–E)** Different letters represent p < 0.05, one-way ANOVA followed by Tukey’s HSD test.

### Infection increases systemic cytokinin response and levels

To understand how haustoria are systemically repressed in *Phtheirospermum*, we undertook a genome-wide RNAseq analysis in *Phtheirospermum* shoots at 10 dpi and found several hundred genes differentially expressed compared to control uninfected plants (Supplemental Fig S1C-E). These included genes related to DNA replication, signal transduction, cell wall modification and response to biotic stimuli (Supplemental Fig S1E). Given that hormones are important for haustoria formation (15, 23), we analysed previously published transcriptomes of infecting roots (72 hpi, Kokla et al 2022) and our infecting shoot datasets for differentially expressed genes related to auxin, cytokinin, brassinosteroid and gibberellic acid. Genes related to brassinosteroids (*BRI1-ASSOCIATED RECEPTOR KINASE* (*PjBAK1*), *BR INSENSITIVE 1* (*PjBRI1*), *BRASSINOSTEROID-INSENSITIVE 2* (*PjBIN2*), *BRASSINAZOLE RESISTANT-1*(*PjBZR1*)) and gibberellic acid *(BETA HLH PROTEIN 93* (*PjbHLH93*), *REPRESSOR OF GA* (*PjRGA1*), *GIBBERELLIN 20-OXIDASE 1* (*PjGA20ox1*), *GA INSENSITIVE DWARF1* (*PjGID1*)) did not show a clear trend of differential expression in either shoot or root, whereas auxin-related genes (*INDOLE-3 ACETIC ACID 14* (*PjIAA14)*, *YUCCA3* (PjYUC3), *LIKE AUXIN RESISTANT* (*PjLAX1*), *PIN-FORMED1* (*PjPIN1*)) were differentially expressed only locally at the site of infection (Fig. 2A, Supplemental Fig S2A). However, cytokinin-related genes *CYTOKININ OXIDASE 3* (*PjCKX3*), *ISOPENTENYLTRANSFERASE 1* (*PjIPT1a*) and *RESPONSE REGULATORs 5b* and *9* (*PjRR5b*, *PjRR9*) were all upregulated in infecting roots, and RRs were also upregulated in shoots of infecting plants (Fig. 2A). This upregulation of cytokinin response at the *Phtheirospermum* infection site occurred for many cytokinin-related genes, already by 12 hours post infection (hpi) for genes like *PjCKX3* and the cytokinin transporters *PURINE PERMEASEs PjPUP1* and *PjPUP3* (Fig. 2B-C, Supplemental Fig. S2B). In infected *Arabidopsis*, little cytokinin-related gene induction was observed by 72 hpi, perhaps in part due to the early sampling points before xylem bridge formation (Fig. 2B). Next, we confirmed the increase in cytokinin signaling during infection using the *pTCSn* cytokinin responsive reporter (24). *pTCSn* was induced by four days after infection in *Phtheirospermum* and *Arabidopsis* at the haustorium site and in the root above haustoria (Fig. 2D). An increase in *pTCSn* signal was also observed in the hypocotyl vasculature of *Arabidopsis* infected at 7 dpi and in flower buds of infected plants at 30 dpi, but not in *Phtheirospermum* roots below the haustoria (Supplemental Fig. S2C). At 10 dpi, roots of infecting *Phtheirospermum* had significantly higher levels of the cytokinin species tZ, tZR, cZ and cZR compared to non-infecting roots (Fig. 3A), while roots of infected *Arabidopsis* showed significantly higher levels of tZ, tZR and iP compared to uninfected controls (Fig. 3B). We then measured the levels of cytokinin in roots and shoots of *Phtheirospermum* in the split-root setup. tZ and tZR levels increased in *Phtheirospermum* infecting roots 10 days after infection, but not in distant non-infecting roots (Fig. 3C, Supplemental Fig. S3A). tZ levels increased in shoots of infecting *Phtheirospermum*, either when one or both sides were infected, while tZR levels only slightly increased in shoots when just one side was infected (Fig. 3C, Supplemental Fig. S3A). Gene expression of the cytokinin-related genes *PjRR5* and *PjHK3* also increased in both roots and shoots when one side of the root system was infected (Supplemental Fig. S3B). The cytokinin biosynthesis homolog *PjIPT1* was upregulated in root RNAseq datasets by infection so we tested its expression by qPCR in shoots and found that it was not significantly increased at 7 dpi in infecting shoots, opposite to that of *PjRR5b* and *9* (Fig. 3D). Together, these results suggest an induction of cytokinin production and response at both the site of local infection and in shoot tissues, consistent with the movement of cytokinins from root to shoot in *Phtheirospermum*.

**Figure 2:**
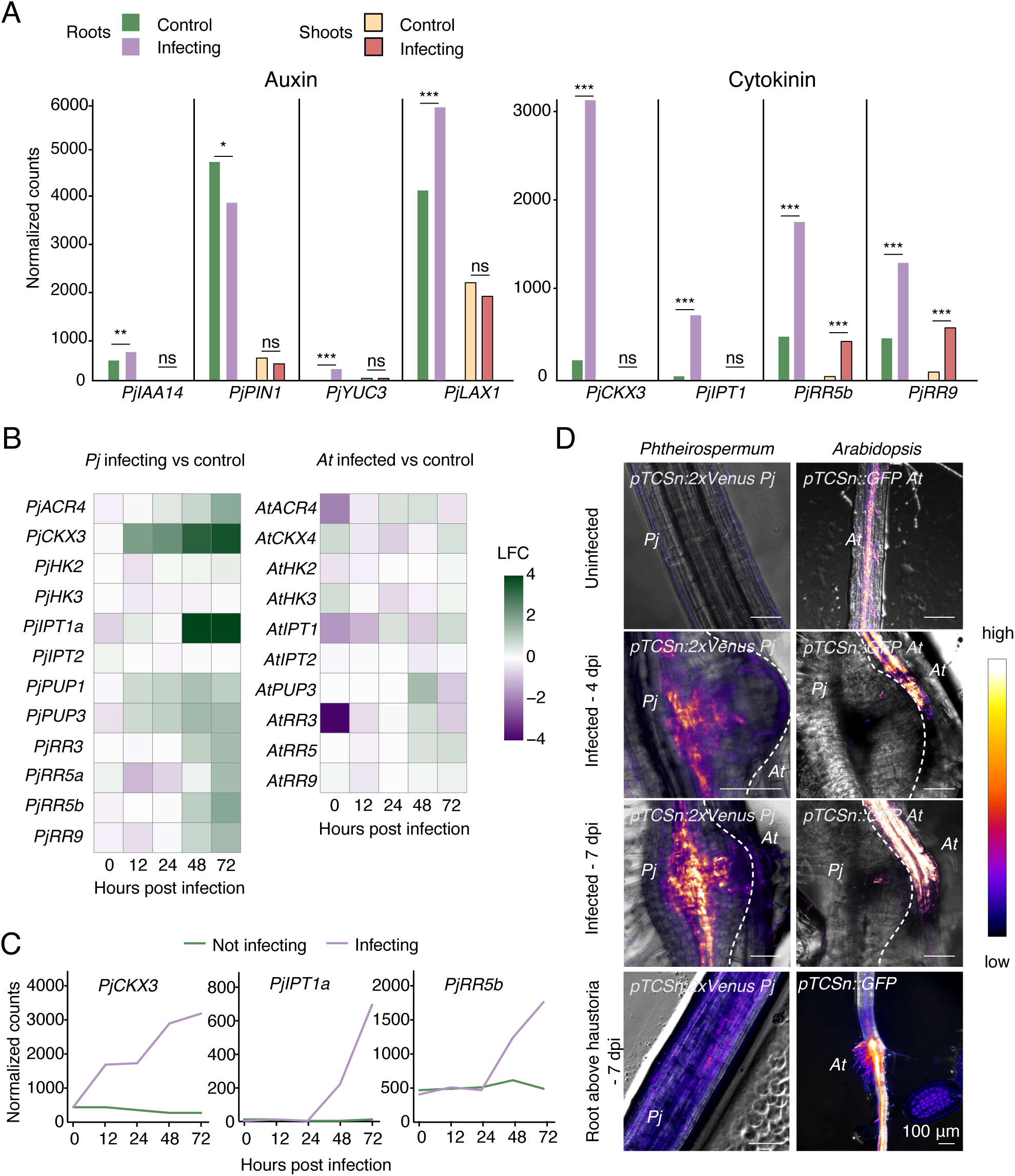
Infection induces systemic transcriptional changes in *Phtheirospermum*. **A)** Normalized counts of auxin and cytokinin-related genes in *Phtheirospermum* infecting or control roots at 72 hours post infection (hpi), and infecting or control shoots at 10 days post infection (n = 3 libraries, Wald test with Benjamini-Hochberg correction, * p<0.05, ** p<0.01, *** p<0.001, ns = not significant) **B)** Normalized counts of *PjCKX3*, *PjIPT1a* and *PjRR5b* in *Phtheirospermum* infecting or control roots at 0, 12, 24, 48 or 72 hpi. **C)** Images of fluorescent *pTCSn* cytokinin reporters in *Arabidopsis* (*At*) or *Phtheirospermum* (*Pj*) for uninfected controls, 4 or 7 dpi haustoria and root above haustoria. Scale bars 100 μm. **D)** Heatmaps of cytokinin-related genes at 0, 12, 24, 48 or 72 hpi in water infect versus water control RNAseq libraries, in *Phtheirospermum* and *Arabidopsis* roots. LFC= log2 fold change.

**Figure 3:**
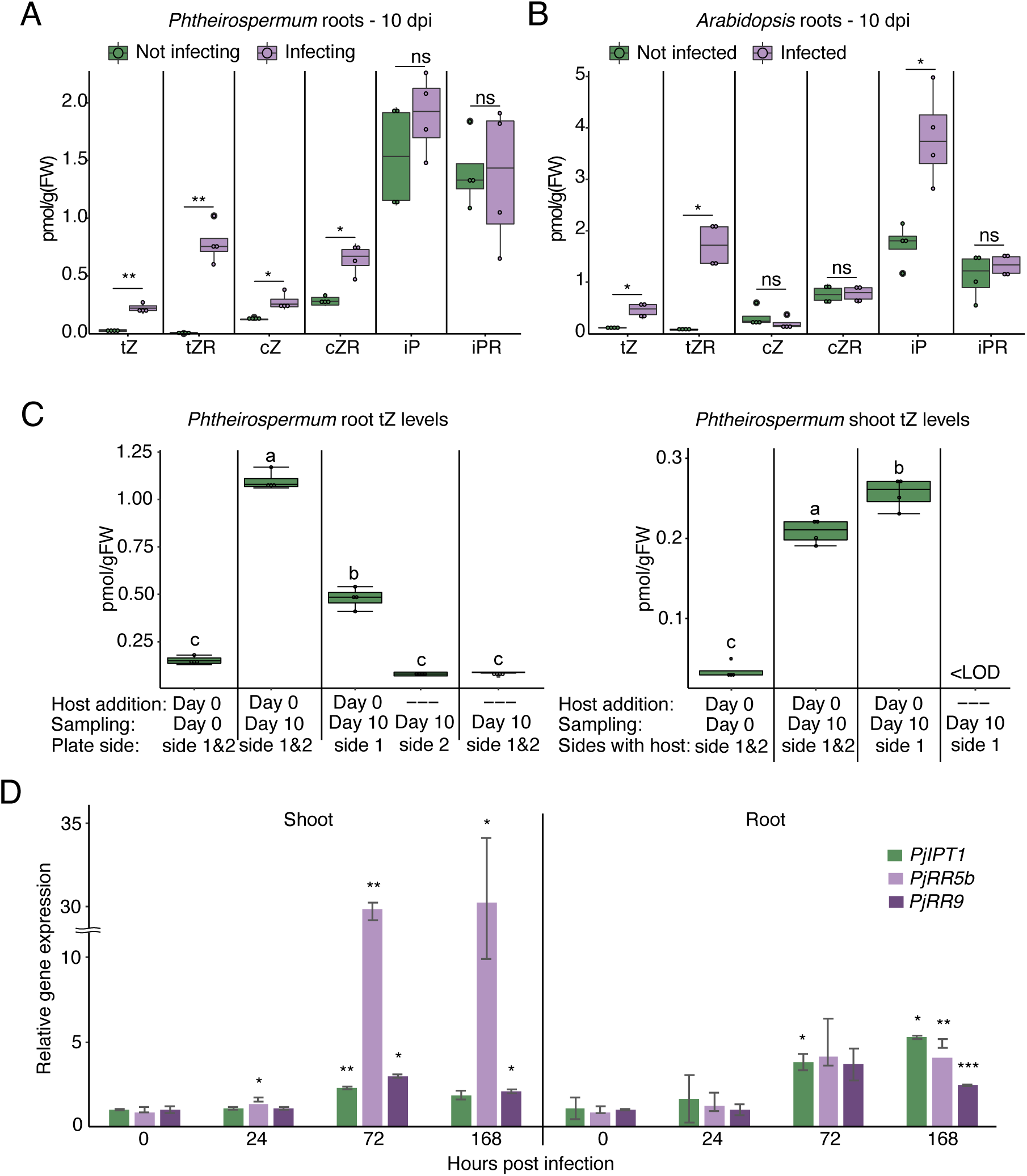
Cytokinin signalling increases systemically in *Phtheirospermum* after infection. **A)** Hormonal quantification of cytokinins in *Phtheirospermum* roots infecting or not infecting *Arabidopsis* at 10 days post infection (dpi). (n = 4 replicates, Student t-test with Benjamini-Hochberg correction, * p<0.05, ** p<0.01, ns= not significant) **B)** Hormonal quantification of cytokinins in *Arabidopsis* roots infected or not infected by *Phtheirospermum* at 10 dpi. (n = 4 replicates, Student t-test with Benjamini-Hochberg correction, * p<0.05, ns= not significant) **C)** Quantification of tZ levels in *Phtheirospermum* roots and shoots in a split-root experimental setup. (n = 4 replicates, one-way ANOVA followed by Tukey’s HSD test) **D)** qRT-PCR gene expression quantification of cytokinin-related genes in *Phtheirospermum* shoots and roots at 0, 24, 72, 168 hours post infection (hpi), normalized to 0 hpi. (n = 2 replicates, Student’s t-test* p<0.05, ** p<0.01, *** p<0.001).

### Cytokinin is a local inhibitor of haustoria development

An increase in cytokinin levels during plant parasitism is associated with host root growth (7), but the role of the cytokinin increase in the parasite is unknown. We performed exogenous cytokinin treatments during infection assays. Application of 80 nM BA, 100 μM kinetin and 1 μM trans-zeatin significantly reduced haustoria induction (Fig 4A,B), while inhibiting cytokinin signaling by applying the cytokinin antagonist PI-55 (1 μM) increased haustoria numbers (Fig 4A-B). To test if this cytokinin-mediated haustoria inhibition required a host, we induced pre-haustoria using DMBQ with no host, and found exogenous cytokinins significantly reduced the number of DMBQ-induced pre-haustoria, while PI-55 had no significant effect (Fig. 4C, Supplemental Fig S3C). We infected the *Arabidopsis* cytokinin related mutants *cre1ahk3*, *ckx3ckx5*, *p35S:CKX1*, *arr1,12*, *arrx8*, *ahp6-3* and *ipt161* and observed no significant difference in haustoria numbers compared to wild-type Col-0 control (Supplemental Fig S3D). We then analyzed a transcriptome dataset where *Phtheirospermum* infecting *Arabidopsis* was treated with BA and haustorium tissues were harvested at 0, 12 and 24 hpi (Kokla et al. 2022). More than 1000 genes were differentially expressed in the BA-treated infecting samples compared to the water infecting samples and in BA infecting versus BA control samples (Supplemental Fig. S4A). Co-expression analyses of BA infect compared to water infect datasets identified three different patterns of expression (Supplemental Fig. S4B). Cluster 1, whose gene expression decreased at 12 hpi, had an overrepresentation of genes related to protein processing, sucrose transport and transcription regulation (Supplemental Fig. S4C). Cluster 2, whose gene expression peaked at 12 hpi, had an over representation of genes related to transcriptional and translational processes (Supplemental Fig. S4C). Cluster 3, whose gene expression peaked at 24 hpi, had an over representation of genes related to oxidoreduction processes (Supplemental Fig. S4C). Many of the genes that were highly upregulated during parasitism were downregulated by BA treatment, consistent with BA repressing a haustoria inducing program (Fig. 4D). Furthermore, genes that were BA responsive in *Phtheirospermum* and *Arabidopsis* were also upregulated during infection, consistent with cytokinin response increasing as the haustoria matured (Fig. 4E, Supplemental Fig. S4E). To further investigate the role of cytokinin, we overexpressed the cytokinin degrading *Arabidopsis* CKX3 enzyme in *Phtheirospermum* hairy roots (Supplemental Fig S4F). Transformed hairy roots formed significantly more haustoria on Col-0 hosts compared to non-transgenic hairy roots (Fig 4F), consistent with parasite-derived cytokinins being important for inhibiting haustoria formation.

**Figure 4:**
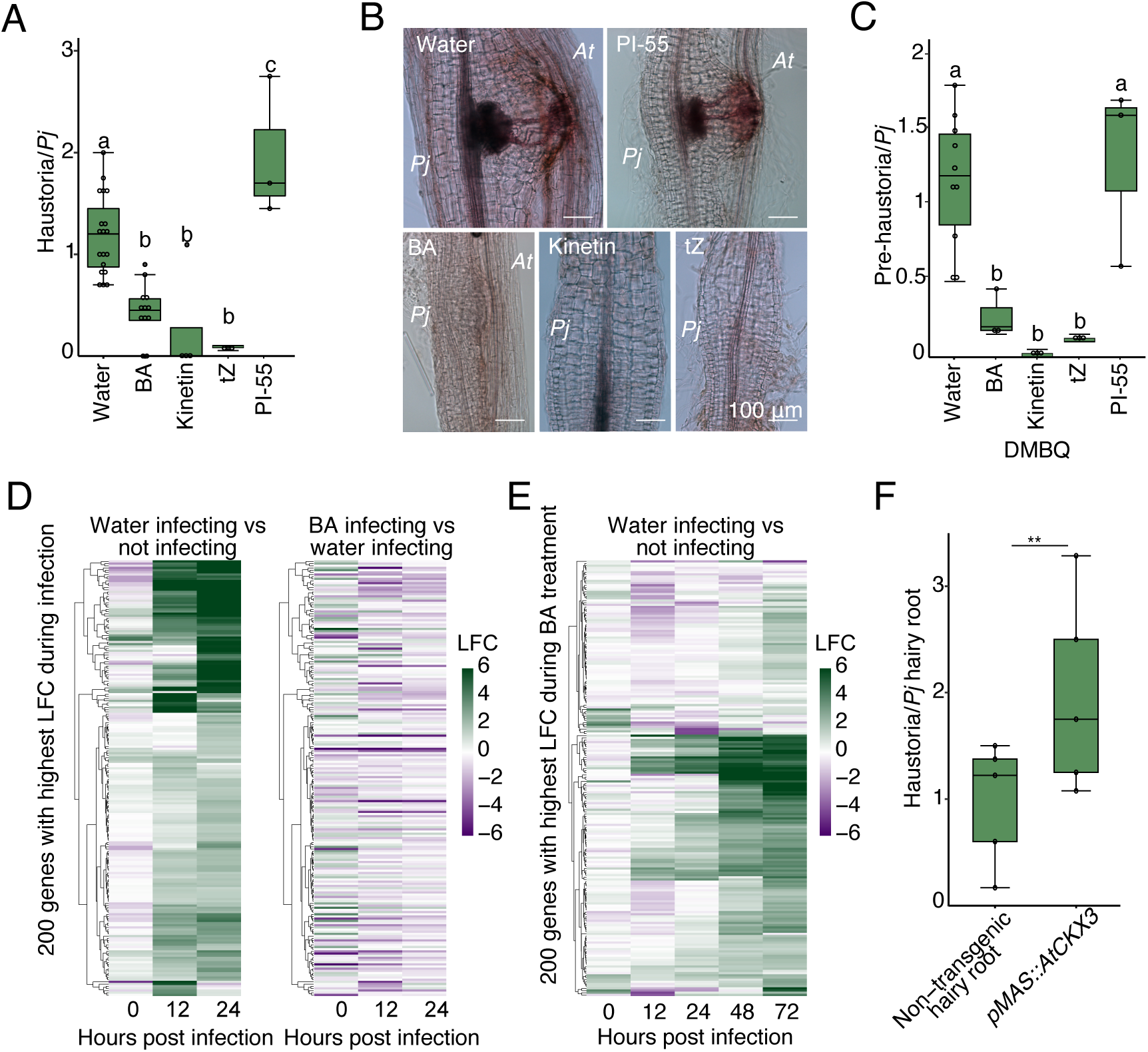
Cytokinin is a local inhibitor of haustoria development. **A)** Average number of haustoria per *Phtheirospermum* seedling in *in vitro* infection assays with 80 nM BA, 100 μM kinetin, 1 μM tZ, 1 μM PI-55 or water control at 7 days post infection (dpi). (n = 3-21 replicates, one-way ANOVA followed by Tukey’s HSD test) **B)** Brightfield images of Safranin-O stained *Phtheirospermum* (*Pj*) haustoria at 7 dpi with chemical treatments. Scale bars 100 μm. **C)** Average number of pre-haustoria per *Phtheirospermum* seedling in *in vitro* haustorium induction assays with 10 μM DMBQ and water, 80 nM BA, 100 μM kinetin, 1 μM tZ or 1 μM PI-55. (n = 3-10 replicates, one-way ANOVA followed by Tukey’s HSD test) **D)** Heatmap of 200 genes with the highest log2 fold change (LFC) during haustoria formation shown over three time points in water infecting versus not infecting and BA infecting versus water infecting *Phtheirospermum* root RNAseq libraries. The water infecting versus water not infecting heatmap was presented in Kokla et al. 2022. **E)** Heatmap of the 200 genes with the highest log2 fold change (LFC) after BA treatment shown over five time points in water infecting versus not infecting *Phtheirospermum* roots RNAseq libraries **F)** Haustoria per *Phtheirospermum* root in infection assay using *Phtheirospermum* with non-transgenic hairy roots or hairy roots overexpressing *AtCKX3*. (n = 5 replicates, Student t-test, ** p<0.01)

### Cytokinin mediates a systemic haustoria repressing signal

We next investigated the role of cytokinin in long distance haustoria repression. When 80 nM BA was applied to one side of the split-root setup, both treated and distant root sides significantly reduced haustoria numbers compared to the control (Fig. 5A), suggesting a systemic repressive role for cytokinin. We then investigated if the local increase in cytokinin biosynthesis and signaling following infection is needed for systemic repression. We treated one side of the split-root setup with 1 μM PI-55 and found PI-55 treatment had no significant effect on haustoria numbers at day 0. However, the distant side infected at 10 days had slightly reduced haustoria numbers but not significantly different to the PI-55 treated side infected at 0 days (Fig. 5B), showing local application of PI-55 partially inhibited the repressive signal in distant roots. Finally, we sought to endogenously modify cytokinin signaling in *Phtheirospermum* by infecting transgenic hairy roots overexpressing *AtCKX3* at day 0, followed by infection of a non-transgenic hairy root on the same plant at day 10. While the control non-transgenic plants showed fewer haustoria on the day 10 root side (Fig. 5C), the transformed plants showed no significant difference in the numbers of haustoria between the day 0 and day 10 infections (Fig. 5C). Taken together, these data indicated that local cytokinin production or response in infecting roots was needed to initiate systemic signaling that regulates future haustoria formation in *Phtheirospermum* (Fig. 5D).

**Figure 5:**
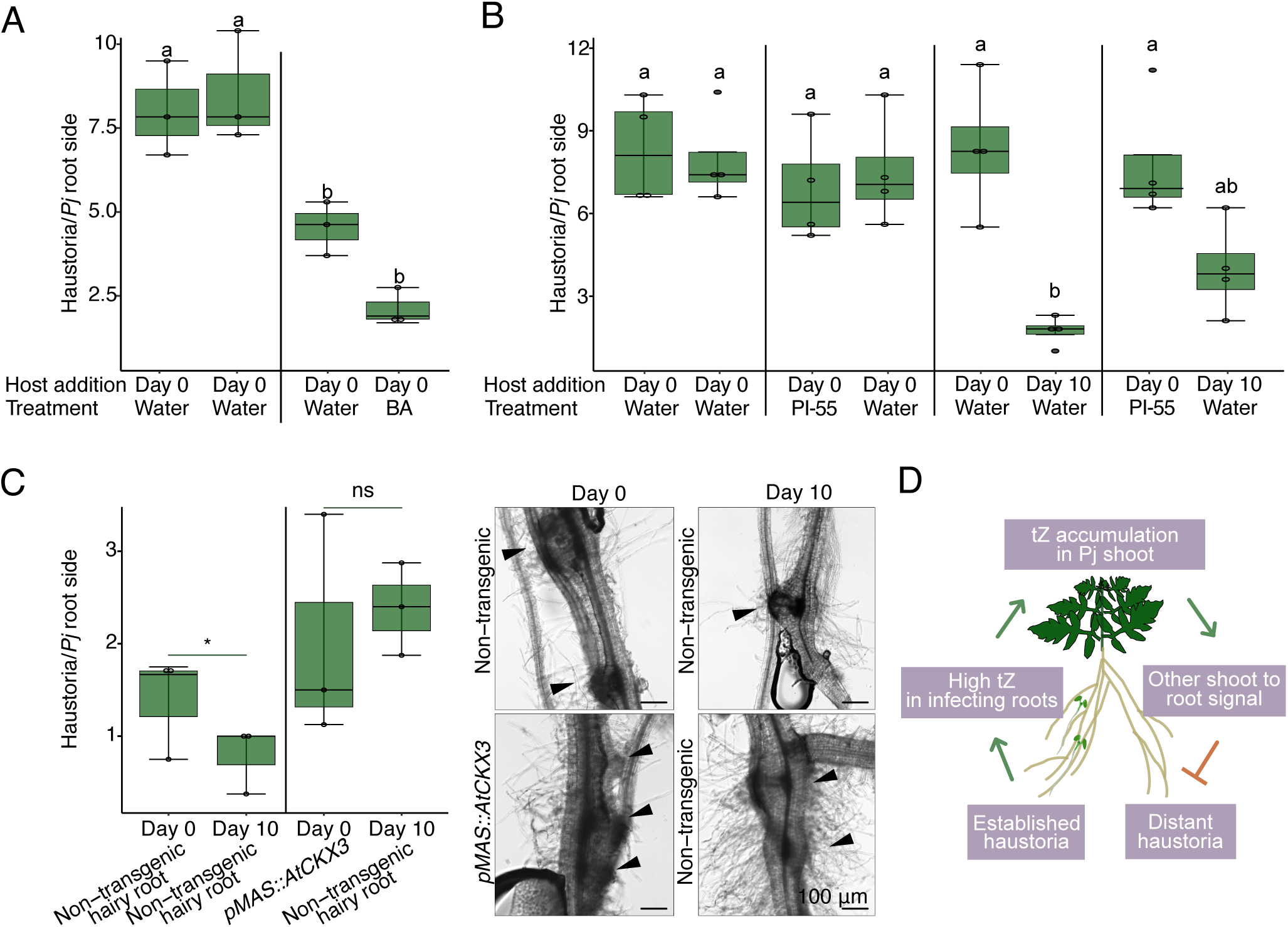
Cytokinin mediates the systemic regulation of haustoria numbers. **A)** Average number of haustoria per *Phtheirospermum* in a split-root setup on water agar or 80 nM BA, with host added on both sides at 0 days post infection (dpi)(n = 3 replicates, one-way ANOVA followed by Tukey’s HSD test) **B)** Average number of haustoria per *Phtheirospermum* in a split-root setup on water agar or 1 μM PI-55, with host added on both sides at 0 dpi or one side at 0 dpi and 10 days later on the other side. (n = 4 replicates, one-way ANOVA followed by Tukey’s HSD test) **C)** Average number of haustoria per *Phtheirospermum* in a split-root setup with non-transgenic hairy roots or hairy roots overexpressing *AtCKX3*, and representative brightfield images. Arrowheads point at haustoria, scale bars 100 μm. (n = 3 replicates, Student’s t-test, * p<0.05 ns = not significant) **D)** Proposed model of the cytokinin-mediated haustoria autoregulation in *Phtheirospermum*.

## Discussion

Plants involved in nutrient-acquiring symbioses with mycorrhiza and nitrogen fixing bacteria regulate their extent of symbiosis (22) and here, we uncover in the facultative parasitic plants *Phtheirospermum japonicum* and *Parentucellia viscosa* a previously undescribed regulatory system whereby existing haustoria control the formation of new haustoria. This system required cytokinin whose increase during parasitism is known from work in *Phtheirospermum*, *Cuscuta* and *Santalum* yet, to date, this increase has only been associated with a developmental response in the host (7, 25, 26). Such parasite-derived cytokinins promote tissue expansion in the host but notably not in the parasite (7), suggesting different roles for these cytokinins between parasite and host. By using a combination of exogenous treatment assays and transgenic approaches, we demonstrated here that parasite-derived cytokinins act as haustoria repressing factors that control local and long-distance haustoria numbers. The identity of the mobile signals mediating systemic repression remain unknown given that our exogenous application assays and transgenic studies focused on the infecting roots. However, trans-zeatin species were highly increased in shoot despite little or late upregulation of cytokinin biosynthesis genes in the shoot, and cytokinin transporters were also updated in infected roots (Fig. 3A,C, Supplemental Fig S2B, Supplemental Fig S3A). Thus, we propose that root-produced cytokinins likely act as a root-to-shoot signaling component associated with systemic suppression, consistent with the known mobility of cytokinins from root to shoot through the xylem in *Arabidopsis* (27). We did not observe an increase in cytokinins in distant roots (Fig. 3C, Supplemental Fig S3A), indicating the shoot-to-root haustoria repressing factor remains unknown.

Notably, in both *Striga hermonthica* and *Phelipanche ramosa*, cytokinins act as haustoria inducing factors (28, 29). These findings contrast with our own data and suggest that cytokinins have different roles in the obligate parasites *Striga* and *Phelipanche* compared to the facultative parasite *Phtheirospermum*, perhaps due to their lifestyle or physiology. *Striga* and *Phelipanche* form haustoria by differentiating the ends of primary root tips, and there, haustoria regulation may occur via a primary root formation or differentiation pathway. In contrast, *Phtheirospermum* form lateral haustoria where divisions in the epidermis, cortex, stele and endodermis are relevant (30). Lateral root initiation is inhibited by cytokinin (31), whereas cytokinins promote primary root differentiation (32) providing a possible explanation for differences between *Striga* and *Phtheirospermum*. Lateral root numbers are also controlled by nutrient-based feedback regulation in plants and a similar situation may occur with haustoria in parasitic plants. Providing low or moderate nitrogen levels promotes lateral root formation, whereas high nitrogen levels suppress lateral root formation (33). In facultative parasitic plants, exposure to a nutrient source like a host might promote haustoria formation, while an abundance of haustoria and hosts might suppress more haustoria, forming parallels to lateral root formation during nitrogen foraging and demonstrating the importance of root plasticity.

Successful symbioses are often characterized by a combination of negative and positive regulators. In legumes, auto-regulation of nodules integrates environmental inputs such as nitrogen with a negative regulation pathway, CLE-SUNN, and a positive regulation pathway, CEP-CRA2, to optimize nodule numbers (34). In the parasitic plant *Phtheirospermum*, several positive signals have been identified, including haustoria inducing factors such as DMBQ, and hormones such as auxin that promotes haustoria initiation, and ethylene that promotes host invasion (11, 15, 16). Recent work in *Phtheirospermum* has found that auxin-related compounds move from shoots to root to promote haustoria maturation (35) and also identified CLE peptides as positive regulators of haustoria formation (36). However, in *Phtheirospermum* and *Striga*, only nitrogen has been identified as a local haustoria suppressing factor (17) and here, we show that cytokinin is repressive, but also extend this previous observation to show that nutrients can also act systemically to regulate haustoria numbers. Such environmental information as well as negative and positive regulators could form parts of long-distance system regulating haustoria numbers. Given the presumed need to balance resource expenditure with resource acquisition during parasitism, we expect there to be additional negative regulators of a haustoria formation in *Phtheirospermum*, *Striga* and other parasitic plants. Discovering these compounds and understanding the prevalence of systemic haustoria regulation and how such a system functions in parasitic plants should be a priority. In addition, deploying and introducing such negative regulators in the host could provide durable resistance to parasites.

## Materials and methods

### Plant materials and growth conditions

*Phtheirospermum japonicum* (Thunb.) Kanitz ecotype Okayama seeds were described previously (Yoshida and Shirasu, 2009). *Parentucellia viscosa* (Pv) seeds used in this study were harvested at Nagoya, Japan in 2023. *Arabidopsis* ecotype Columbia (Col-0) was used as *Arabidopsis* wild-type (WT). *Arabidopsis cre1-12ahk3-3*, *ckx3-1ckx5-2, 35S::CKX1, arr1,12*, *arrx8*, *ahp6-3*, *ipt161*, *pTCSn::GFP* were published previously (24, 37–43). For *in vitro* germination, *Phtheirospermum* and *Arabidopsis* seeds were surface sterilized with 70% (v/v) EtOH for 20 minutes followed by 95%(v/v) EtOH for 5 minutes. The seeds were then placed on ½MS medium (0.8% (w/v) plant agar, 1% (w/v) sucrose, pH 5.8). After overnight stratification in the dark and 4°C, the plants were placed to long day conditions (16-h light:8- h dark and light levels 100 μmol m^−2^ s^−1^) and 25°C. For *Parentucellia*, 〜500 seeds were surface sterilized in 1.5 mL tube by rinsing with 20% solution of commercial bleach for three times, followed by ethanol three times and distilled water for three times. The tube was covered with tin foil and rotated at 16℃ for one week. The seeds were plated on 0.7% agarose/water plate and germinated for two weeks at 16℃ in 12h light:12h dark condition.

### In vitro infection assays with Phtheirospermum

The *in vitro* infection assays were performed following Kokla et al. 2022. Four to five-day-old *Phtheirospermum* seedlings were transferred for three days to nutrient-free 0.8% (w/v) agar medium or 0.8% (w/v) agar medium supplemented by hormone treatments: 0.08 μM 6- Benzylaminopurine (BA), 1 μM PI-55, 1 μM trans-zeatin or 100 μM kinetin. Five-day-old *Arabidopsis* seedlings were aligned next to and roots place in contact with the pre-treated *Phtheirospermum* roots for infection assays. Haustoria numbers were measured at seven days post infection using a Zeiss Axioscope A1 microscope. For pre-haustorium assays, seedlings after the three-day starvation/hormone treatment were transferred to 0.8% (w/v) agar medium containing 10 μM DMBQ (Sigma-Aldrich) with or without hormonal treatment. Pre-haustoria were counted at seven days.

### Split-root infection assays with *Phtheirospermum* and *Parentucellia*

1 month old *Phtheirospermum* were transferred on Gosselin™ polystyrene round petri plates (split-plate) (Fisher Scientific) with nutrient-free 0.8% (w/v) agar medium on both sides or 0.8% (w/v) agar medium supplemented by 10.3 mM NH_4_NO_3_, 0.08 μM BA, 1 μM ABA or 1 μM PI-55 on one side. The root system separated in the 2 sides of the split-plate so they are not in contact with each other. Four days later two 7 days old *Arabidopsis* (Col-0) hosts were added in alignment with the roots of *Phtheirospermum* on either both sides or one of them. In the non-host root sides, the hosts were added 10 days later (10 dpi). The measurements of haustoria numbers were taken at 7 days after host addition. For split root assays with *Parentucellia*, seedlings were incubated after germination in the same growth condition as for the *Phtheirospermum* split root assay until the root length reached to 2-3 cm (〜1.5 months). The assay was then performed as described for *Phtheirospermum*.

### Cloning of pTCSn:2xVenus-NLS for expression in Phtheirospermum

The cloning was based on the Greengate method following the standard protocols (Lampropoulos et al. 2013). The primers used for GreenGate cloning are listed in Supplemental Table S3. Digestion and ligation reactions were performed using the BsaI- HFv2 (NEB #R3733) and T4 DNA Ligase (NEB #M0202) enzymes respectively. The *pTCSn* promoter fragment was cloned from a previously published vector (7) using the CloneAmp™ HiFi PCR Premix (TakaraBio) and inserted into the entry vector *pGGA000* (Addgene plasmid #48856) to create the *pGGA-pTCSn* vector. The ligated plasmid was amplified in *E. coli* DH5α and confirmed by Sanger sequencing. The binary vector assembly was performed using *pGGA-pTCSn, pGGB003* (Addgene plasmid #48821*), pGGC-2xVenus-NLS* (14), *pGGD002* (Addgene plasmid #48834), *pGGE-tMAS* (17), *pGGF- DsRed* (17) and *pGGZ001* (Addgene plasmid #48868). The final plasmid was co-transformed in electrocompetent *Agrobacterium rhizogenes* AR1193 with the pSoup plasmid (Addgene plasmid #165419), and the bacteria cultured in LB broth with 50 ug/ml spectinomycin and 50 ug/ml rifampicin.

### Phtheirospermum transformation

Transformation was performed as described previously (Ishida et al., 2011). Briefly, five-day-old *Phtheirospermum* seedlings were sonicated for 10 to 15 seconds followed by vacuum infiltration for 5 minutes with suspension of *Agrobacterium rhizogenes* strain AR1193 carrying the *Arabidopsis CKX3* gene overexpressing construct, *pMAS::AtCKX3:tMas*, previously described (Spallek et al., 2017), or *pTCSn::2xVenus-NLS:tMas*. The seedlings were then transferred on co-cultivation media (Gamborg’s B5 medium, 0.8% agar, 1% sucrose, 450 μM acetosyringone) first at 20°C for 2 days in dark conditions, then at 25°C under long-day conditions for ∼ 1 month with 300 μg/ml cefotaxime. Transgenic hairy roots were infected as described above. Both transgenic and non-transgenic hairy roots showed reduced levels of infection compared to non-hairy roots, likely due to transformation conditions slightly reducing plant vigor.

### Histological staining, microscopy and confocal imaging

For visualization of xylem bridges, dissected *Phtheirospermum* roots were fixed in ethanol-acetic acid and stained with Safranin-O solution (0.1%) followed by clearing with chloral hydrate for two to three days before microscopic observation with a Zeiss Axioscope A1 microscope as described previously (Cui et al., 2016). A Leica M205 FA stereo microscope was used with RFP filter for the selection of transformed *Phtheirospermum* hairy roots. The *TCSn* fluorescent reporters was visualized in *Arabidopsis* root-hypocotyl junction, hypocotyl and flowers using a Leica M205 FA stereo microscope with GFP filter. *Arabidopsis* and *Phtheirospermum TCSn* roots were visualized on a Zeiss LSM780 laser scanning confocal microscope with 514 nm excitation, 2.5% laser power, 518 to 624 nm emission.

### Sample preparation for RNAseq

Sample preparation for the root infection time course in control (water) conditions and following BA treatment is described in Kokla et al. 2022. For the preparation of the *Phtheirospermum* shoot sequencing, four *Phtheirospermum* seedlings of four to five days old per sample were transferred to nutrient-free 0.8% (w/v) agar medium for 3 days prior to infection with *Arabidopsis* Col-0. As a control, four *Phtheirospermum* seedlings remained without the *Arabidopsis* host. The shoot of the *Phtheirospermum* seedlings were collected at 10dpi. These experiments were replicated three times. RNA extraction was performed using the ROTI®Prep RNA MINI (Roth) kit following the manufacturer’s instructions. The isolation of mRNA and library preparation were performed using NEBNext® Poly(A) mRNA Magnetic Isolation Module (#E7490), NEBNext® Ultra™ RNA Library Prep Kit for Illumina® (# E7530L), NEBNext® Multiplex Oligos for Illumina® (#E7600) following the manufacturer’s instructions. The libraries were then sequenced using paired-end sequencing with an Illumina NovaSeq 6000.

### Bioinformatic analysis

Bioinformatic analysis of the water and BA sequencing data have been previously described (Kokla et al., 2022). The same method was followed for the shoot sequencing data. Briefly, the adapter and low-quality sequences were removed using the fastp software with default parameters (Chen et al., 2018). The quality-filtered reads were mapped to both the *Phtheirospermum* genome (Cui et al., 2020) and *Arabidopsis* genome (TAIR10) using STAR (Dobin et al., 2013) and were separated based on mapping to *Phtheirospermum* and *Arabidopsis* reads. The separated reads were then re-mapped to their respective genomes. The read count was calculated using FeatureCounts (Liao et al., 2014). The differential expression analysis was performed using Deseq2 (Love et al., 2014) (Table S1). The gene expression clustering was performed using the Mfuzz software (Futschik et al., 2009).

Custom annotations of the *Phtheirospermum* predicted proteins (Cui et al., 2020) were estimated using InterProScan (Blum et al., 2020) and used for the gene ontology analysis that was performed using the topGO software (Alexa et al., 2016). Cytokinin related genes from *Arabidopsis* were blasted against the *Phtheirospermum* genome (Cui et al., 2020) using the tBLASTp and tBLASTn algorithms; these genes can be found in Table S2. The 200 genes with highest expression were identified by selecting the genes with LFC>1.5 in the *Phtheirospermum* water infecting vs not infecting RNAseq libraries for the 12 and 24 hpi time points, *Phtheirospermum* BA vs DMSO not infecting RNAseq libraries for the 0 hpi time point, and *Arabidopsis* BA vs DMSO not infected RNAseq libraries for the 24 hpi time point (Table S2).

### qRT-PCR

For the split root setup, the root sides or shoots of three plants were collected for each replicate. For the infection time course, the roots or shoots of 5 infecting or control *Phtheirospermum* seedlings were harvested at 0, 24, 72 and 168 hours post infection and frozen in liquid nitrogen. For the *AtCKX3* transformed hairy roots, four or five fluorescent or non fluorescent (control) hairy roots were collected for each replicate. RNA extraction was performed using the ROTI®Prep RNA MINI (Roth) kit following the manufacturer’s instructions. cDNA synthesis was performed using Maxima First Strand cDNA Synthesis Kit for RT-qPCR (Thermo Scientific™) following the manufacturer’s instructions. *PjPP2A* (Ishida et al., 2016) was used as an internal control. qPCR was performed with SYBR-Green master mix (Applied Biosystems™). The relative expression was calculated using the Pfaffl method (Pfaffl, 2001). The primers used for this experiment are listed in Table S3.

### Hormonal quantification

For hormonal quantifications of roots in *Phtheirospermum* and *Arabidopsis*, *Phtheirospermum* seedlings were grown for five days before transferring to water agar for three days. *Arabidopsis* Col-0 was then placed next to the *Phtheirospermum* seedlings and left for 10 days. *Phtheirospermum* with no host were used as a control. Four to five *Phtheirospermum* or *Arabidopsis* whole seedlings were then harvested per sample with care taken to separate samples under a microscope to mimimize sample mixing. For quantification on the split plate setup, ∼ 1month *Phtheirospermum* seedlings were placed on nutrient-free 0.8% (w/v) agar medium or 0.8% (w/v) agar medium on petri dishes with a plastic separator in the middle (split-plate) for seven days. *Arabidopsis* Col-0 was placed next to the *Phtheirospermum* roots on both or one of the split roots sides and left for 10 days. *Phtheirospermum* at 0 dpi were used as control. The root system of three *Phtheirospermum* plants per sample were collected separate for each root side at 0 dpi and 10 dpi with care taken to separate host and parasite under a microscope to mimimize sample mixing. The samples were crushed to powder using liquid nitrogen with mortar and pestle. Samples were extracted, purified and analyzed according a previously published method (Šimura et al., 2018). Briefly, approx. 20 mg of frozen material per sample was homogenized and extracted in 1 mL of ice-cold 50% aqueous acetonitrile (v/v) with the mixture of ^13^C- or deuterium-labelled internal standards using a bead mill (27 hz, 10 min, 4°C; MixerMill, Retsch GmbH, Haan, Germany) and sonicator (3 min, 4°C; Ultrasonic bath P 310 H, Elma, Germany). The samples were then centrifuged (14 000 RPM, 15 min, 4°C) and the supernatant was purified according to the following procedure. A solid-phase extraction column Oasis HLB (30 mg 1 cc, Waters Inc., Milford, MA, USA) was conditioned with 1ml of 100% methanol and 1ml of deionized water (Milli-Q, Merck Millipore, Burlington, MA, USA) followed by sample loading on the SPE column. The flow-through and elution fraction was collected 1ml 30% aqueous acetonitrile (v/v). The samples were then dried using speed vac (SpeedVac SPD111V, Thermo Scientific, Waltham, MA, USA) and dissolved in 40 µL of 30% acetonitrile (v/v) in insert-equipped vials. Mass spectrometry analysis for the detection of targeted compounds was performed by an UHPLC-ESI-MS/MS system comprising of a 1290 Infinity Binary LC System coupled to a 6490 Triple Quad LC/MS System with Jet Stream and Dual Ion Funnel technologies (Agilent Technologies, Santa Clara, CA, USA). The quantification was carried out in Agilent MassHunter Workstation Software Quantitative (Agilent Technologies, Santa Clara, CA, USA). Hormonal quantification values are provided in Table S4.

### Accession numbers

Sequence data are available at the Gene Expression Omnibus (http://www.ncbi.nlm.nih.gov/geo/) under accession numbers GSE177484 and GSE253722. Sequence data of the *Phtheirospermum* genes studied in this article are available in GenBank (http://www.ncbi.nlm.nih.gov/genbank/) under the accession numbers provided in TableS5.

## Acknowledgements

AK, ML and CWM were supported by a Wallenberg Academy Fellowship (2016-0274) and an ERC starting grant (GRASP- 805094). KL and JS were supported by grants from the Swedish Research Council, the Swedish Governmental Agency for Innovation Systems and the Knut and Alice Wallenberg Foundation. MH, NN and YT were supported by JSPS Kakenhi (21H04775) and JST CREST (JPMJCR1924). The authors acknowledge support from the Uppsala Multidisciplinary Center for Advanced Computational Science for assistance with access to the UPPMAX computational infrastructure. We also thank the Swedish Metabolomics Centre for access to instrumentation.

## Author contributions

AK, ML and CWM conceived the experiments. AK, ML, MH, NN and JS performed the experiments. KL, YT and CWM supervised the experiments. AK, ML and CWM wrote the paper. All authors edited and revised the final paper.

## Supplemental data

Supplemental Figure S1

Supplemental Figure S2

Supplemental Figure S3

Supplemental Figure S4

Supplemental Table S1: Deseq2 results of *P. japonicum* shoot sequencing

Supplemental Table S2: Genes included in heatmaps

Supplemental Table S3: List of primers used

Supplemental Table S4: Cytokinin hormonal quantification values

Supplemental Table S5: Accession numbers of *P. japonicum* and *A. thaliana* genes shown in figures

## References

1. D. L. Nickrent, Parasitic angiosperms: How often and how many? Taxon 69, 5–27 (2020).

2. C. Parker, Observations on the current status of *Orobanche* and *Striga* problems worldwide. Pest Manag Sci 65, 453–459 (2009).

3. J. Rodenburg, M. Demont, S. J. Zwart, L. Bastiaans, Parasitic weed incidence and related economic losses in rice in Africa. Agriculture, Ecosystems and Environment 235, 306–317 (2016).

4. J. Kuijt, The biology of parasitic flowering plants (University of California Press, Berkeley,, 1969), pp. 246 p.

5. J. Tesitel et al., The bright side of parasitic plants: what are they good for? Plant Physiology 185, 1309–1324 (2021).

6. T. J. Barkman et al., Mitochondrial DNA suggests at least 11 origins of parasitism in angiosperms and reveals genomic chimerism in parasitic plants. Bmc Evol Biol 7 (2007).

7. T. Spallek et al., Interspecies hormonal control of host root morphology by parasitic plants. Proceedings of the National Academy of Sciences 114, 5283–5288 (2017).

8. A. Kokla, C. W. Melnyk, Developing a thief: Haustoria formation in parasitic plants. Developmental Biology 442, 53–59 (2018).

9. S. Shahid et al., MicroRNAs from the parasitic plant Cuscuta campestris target host messenger RNAs. Nature 553, 82–85 (2018).

10. M. Chang, D. G. Lynn, D. H. Netzly, L. G. Butler, Chemical Regulation of Distance: Characterization of the First Natural Host Germination Stimulant for Striga asiatica. Journal of the American Chemical Society 108, 7858–7860 (1986).

11. S. Cui et al., Host lignin composition affects haustorium induction in the parasitic plants Phtheirospermum japonicum and Striga hermonthica. New Phytologist 218, 710–723 (2018).

12. V. Goyet et al., Haustorium Inducing Factors for Parasitic Orobanchaceae. Frontiers in Plant Science 10, 1–8 (2019).

13. S. Cui et al., Haustorial hairs are specialized root hairs that support parasitism in the facultative parasitic plant, Phtheirospermum japonicum. Plant Physiology 170, 1492–1503 (2016).

14. M. Leso, A. Kokla, M. Feng, C. W. Melnyk, Pectin modifications promote haustoria development in the parasitic plant Phtheirospermum japonicum. Plant Physiology 10.1093/plphys/kiad343 (2023).

15. J. K. Ishida et al., Local Auxin Biosynthesis Mediated by a YUCCA Flavin Monooxygenase Regulates Haustorium Development in the Parasitic Plant Phtheirospermum japonicum. The Plant Cell 28, 1795–1814 (2016).

16. S. Cui et al., Ethylene signaling mediates host invasion by parasitic plants. Science Advances 6 (2020).

17. A. Kokla et al., Nitrogen represses haustoria formation through abscisic acid in the parasitic plant Phtheirospermum japonicum. Nature Communications 13 (2022).

18. J. J. Kieber, G. E. Schaller, Cytokinin signaling in plant development. Development 145 (2018).

19. A. Greifenhagen et al., The Phtheirospermum japonicum isopentenyltransferase PjIPT1a regulates host cytokinin responses in Arabidopsis. New Phytologist 232, 1582–1590 (2021).

20. S. Siddique et al., A parasitic nematode releases cytokinin that controls cell division and orchestrates feeding site formation in host plants. Proceedings of the National Academy of Sciences of the United States of America 112, 12669–12674 (2015).

21. V. Mortier, E. De Wever, M. Vuylsteke, M. Holsters, S. Goormachtig, Nodule numbers are governed by interaction between CLE peptides and cytokinin signaling. Plant Journal 70, 367–376 (2012).

22. C. L. Wang, J. B. Reid, E. Foo, The Art of Self-Control - Autoregulation of Plant-Microbe Symbioses. Frontiers in Plant Science 9 (2018).

23. T. Wakatake, S. Ogawa, S. Yoshida, K. Shirasu, Auxin transport network underlies xylem bridge formation between the hemi-parasitic plant Phtheirospermum japonicum and host Arabidopsis. Development 10.1242/dev.160788 (2020).

24. E. Zurcher et al., A Robust and Sensitive Synthetic Sensor to Monitor the Transcriptional Output of the Cytokinin Signaling Network in Planta. Plant Physiol 161, 1066–1075 (2013).

25. X. H. Zhang et al., Endogenous hormone levels and anatomical characters of haustoria in *Santalum album* L. seedlings before and after attachment to the host. Journal of Plant Physiology 169, 859–866 (2012).

26. T. Furuhashi et al., Morphological and plant hormonal changes during parasitization by *Cuscuta japonica* on *Momordica charantia*. J Plant Interact 9, 220–232 (2013).

27. M. Matsumoto-Kitano et al., Cytokinins are central regulators of cambial activity. Proceedings of the National Academy of Sciences of the United States of America 105, 20027–20031 (2008).

28. V. Goyet et al., Haustorium initiation in the obligate parasitic plant Phelipanche ramosa involves a host-exudated cytokinin signal. Journal of Experimental Botany 68, 5539–5552 (2017).

29. N. Aoki, S. K. Cui, S. Yoshida, Cytokinins Induce Prehaustoria Coordinately with Quinone Signals in the Parasitic Plant Striga hermonthica. Plant and Cell Physiology 63, 1446–1456 (2022).

30. T. Wakatake, S. Yoshida, K. Shirasu, Induced cell fate transitions at multiple cell layers configure haustorium development in parasitic plants. Development 145, dev164848-dev164848 (2018).

31. L. Laplaze et al., Cytokinins act directly on lateral root founder cells to inhibit root initiation. Plant Cell 19, 3889–3900 (2007).

32. R. Dello Ioio et al., Cytokinins determine root-meristem size by controlling cell differentiation. Current Biology 17, 678–682 (2007).

33. R. F. H. Giehl, N. von Wirén, Root Nutrient Foraging. Plant Physiology 166, 509–517 (2014).

34. C. Laffont et al., Independent Regulation of Symbiotic Nodulation by the SUNN Negative and CRA2 Positive Systemic Pathways. Plant Physiology 180, 559–570 (2019).

35. P. T. Serivichyaswat et al., High temperature perception in leaves promotes vascular regeneration and graft formation in distant tissues. Development 149 (2022).

36. A. Greifenhagen et al., The peptide hormone Pj CLE1 stimulates haustorium formation in the parasitic plant Phtheirospermum japonicum. Proceedings of the National Academy of Sciences 121(**42**) (2024).

37. M. Higuchi et al., In planta functions of the Arabidopsis cytokinin receptor family. Proceedings of the National Academy of Sciences of the United States of America 101, 8821–8826 (2004).

38. I. Bartrina, E. Otto, M. Strnad, T. Werner, T. Schmülling, Cytokinin Regulates the Activity of Reproductive Meristems, Flower Organ Size, Ovule Formation, and Thus Seed Yield in. Plant Cell 23, 69–80 (2011).

39. T. Werner et al., Root-Specific Reduction of Cytokinin Causes Enhanced Root Growth, Drought Tolerance, and Leaf Mineral Enrichment in *Arabidopsis* and Tobacco. Plant Cell 22, 3905–3920 (2010).

40. M. G. Mason et al., Multiple type-B response regulators mediate cytokinin signal transduction in Arabidopsis. Plant Cell 17, 3007–3018 (2005).

41. A. P. Mähönen et al., Cytokinin signaling and its inhibitor AHP6 regulate cell fate during vascular development. Science 311, 94–98 (2006).

42. E. E. van der Graaff, C. A. Auer, P. J. J. Hooykaas, Altered development of Arabidopsis thaliana carrying the Agrobacterium tumefaciens gene is partially due to ethylene effects. Plant Growth Regulation 34, 305–315 (2001).

43. W. J. Zhang, J. P. C. To, C. Y. Cheng, G. E. Schaller, J. J. Kieber, Type-A response regulators are required for proper root apical meristem function through post-transcriptional regulation of PIN auxin efflux carriers. Plant Journal 68, 1–10 (2011).

