## Supplemental Figures 1-4 for "A long-distance inhibitory system regulates haustoria numbers in parasitic plants"

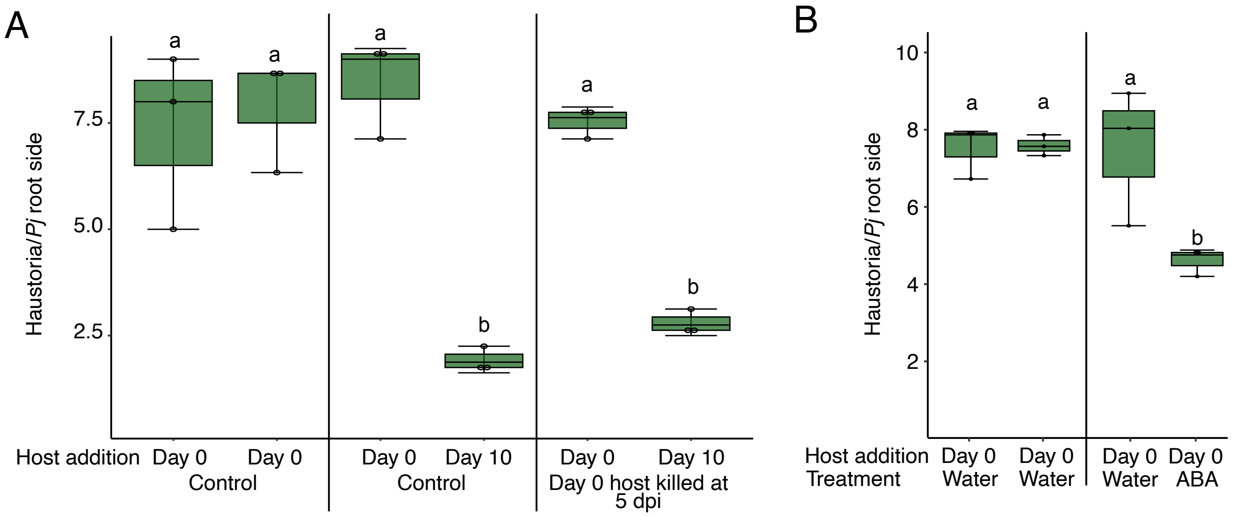


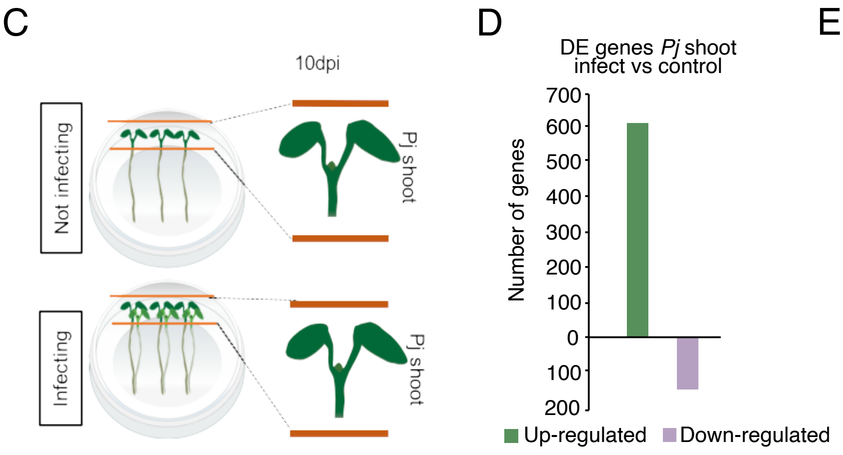


E

| GO term | | p-value |
|---|---|---|
| GO:0006260 | DNA replication | 7.20E-10 |
| GO:0000160 | phosphorelay signal transduction | 3.30E-05 |
| GO:0046274 | lignin catabolic process | 8.20E-05 |
| GO:0046080 | dUTP metabolic process | 0.0007 |
| GO:0009607 | response to biotic stimulus | 0.0019 |
| GO:0044264 | cellular polysaccharide metabolism | 0.0037 |
| GO:0006633 | fatty acid biosynthetic process | 0.0039 |
| GO:0007018 | microtubule-based movement | 0.0068 |
| GO:0006952 | defence response | 0.0098 |
| GO:0071555 | cell wall organization | 0.0122 |

**Figure S1: ABA does not regulate haustoria numbers systemically**

**A)** Average number of haustoria per *Phtheirospermum* in a split-root setup on water agar with host added on both sides at 0 days post infection (day 0)(dpi), or on one side at 0 dpi and other side at 10 dpi. In control samples, hosts were left alive throughout infection, or the host added at day 0 was killed at 5 dpi, before adding the host on the second side (n = 3 replicates, one-way ANOVA followed by Tukey's HSD test) **B)** Average number of haustoria per *Phtheirospermum* root side in a split-root setup on water agar or 1 μM ABA with host added on both sides at 0 dpi. (n = 3 replicates, one-way ANOVA followed by Tukey's HSD test) **C)** Drawings showing the experimental setup for the *Phtheirospermum* shoot sequencing. **D)** Number of genes differentially expressed between control and infecting *Phtheirospermum* shoots at 10 dpi. **E)** Gene ontology analysis for the differentially expressed genes between control and infecting *Phtheirospermum* shoots at 10 dpi (n = 3 libraries, Wald test with Benjamini-Hochberg correction, $p<0.05$).

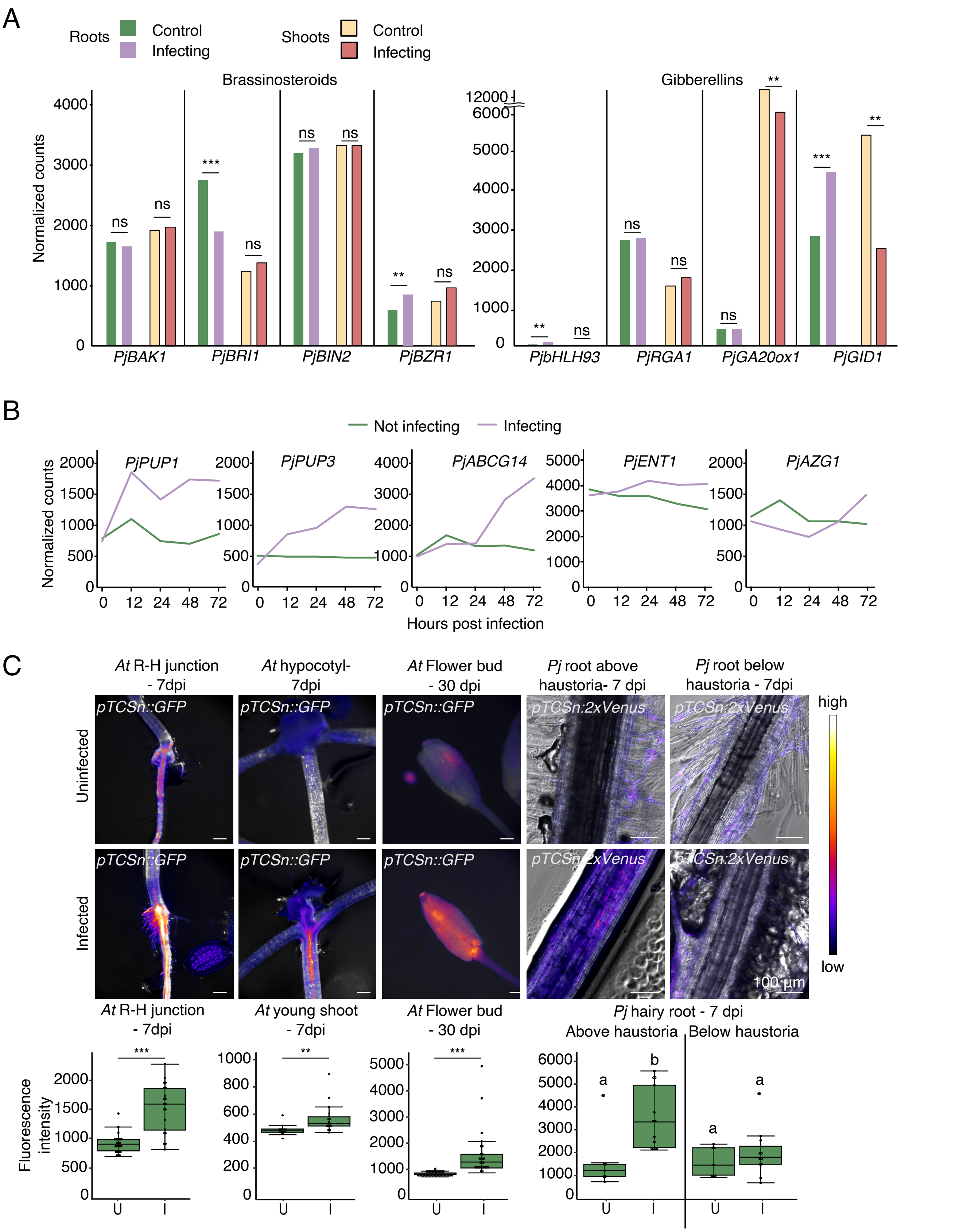


**Figure S2: Brassinosteroid and gibberellins signalling are not systemic in *Phtheirospermum* following infection**

**A)** Normalized counts of brassinosteroid and gibberellin-related genes in *Phtheirospermum* infecting or control roots at 72 hours post infection (hpi), and infecting or control shoots at 10 days post infection (dpi). (n = 3 libraries, Wald test with Benjamini-Hochberg correction, ** p<0.01, *** p<0.001, ns = not significant) **B)** Normalized counts of *PjPUP1*, *PjIPUP3* and *PjABCG14*, *PjENT1* and *PjAZG1* in *Phtheirospermum* infecting or control roots at 0, 12, 24, 48 or 72 hpi. **C)** Images and quantifications of fluorescent *pTCSn* cytokinin reporters for 7 dpi *Arabidopsis* root-hypocotyl (R-H) junctions and shoots, 1 month post infection *Arabidopsis* flower buds and 7 dpi *Phtheirospermum* hairy roots above (upper root) and below haustorium site (lower root). Scale bars 100 µm. *At* = *Arabidopsis, Pj* = *Phtheirospermum*, U = uninfected, I = infected. (for *Arabidopsis*: n = 20-41 images, ** p<0.01, *** p<0.001, Student t-test; for *Phtheirospermum*: n = 7-13 images, one-way ANOVA followed by Tukey's HSD test).

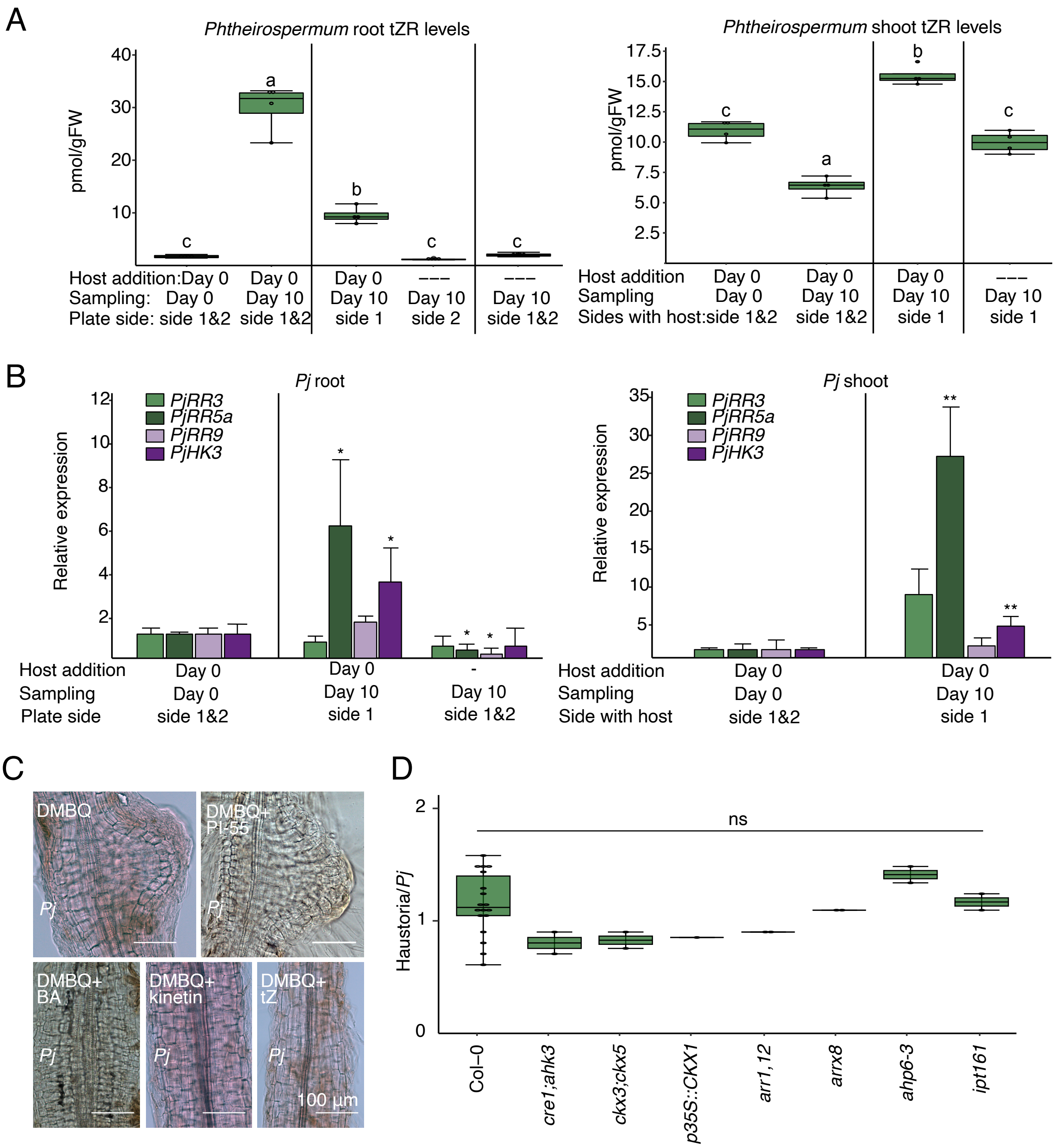


**Figure S3: Host cytokinin response does not influence haustoria induction**

**A)** Quantification of tZR levels in *Phtheirospermum* roots and shoots in a split-root experimental setup. (n = 4 replicates, one-way ANOVA followed by Tukey's HSD test) **B)** qRT-PCR gene expression quantification of cytokinin-related genes in *Phtheirospermum* roots and shoots in a split-root setup. (n = 3 replicates, Student's t-test, * p<0.05, ** p<0.01). **C)** Brightfield images of *Phtheirospermum* (Pj) pre-haustoria at 7 days post infection (dpi) treated with DMBQ, cytokinins or PI-55. Scale bars 100 µm. **D)** Average number of haustoria at 7 dpi per *Phtheirospermum* seedling infecting *Arabidopsis cre1;ahk3*, *ckx3;ckx5*, *p35S::CKX1, aarr1,12, arrx8, ahp6-3* and *ipt161* mutants, or Col-0 control. (n = 2-18 replicates, one-way ANOVA followed by Tukey's HSD test, ns = not significant)

A

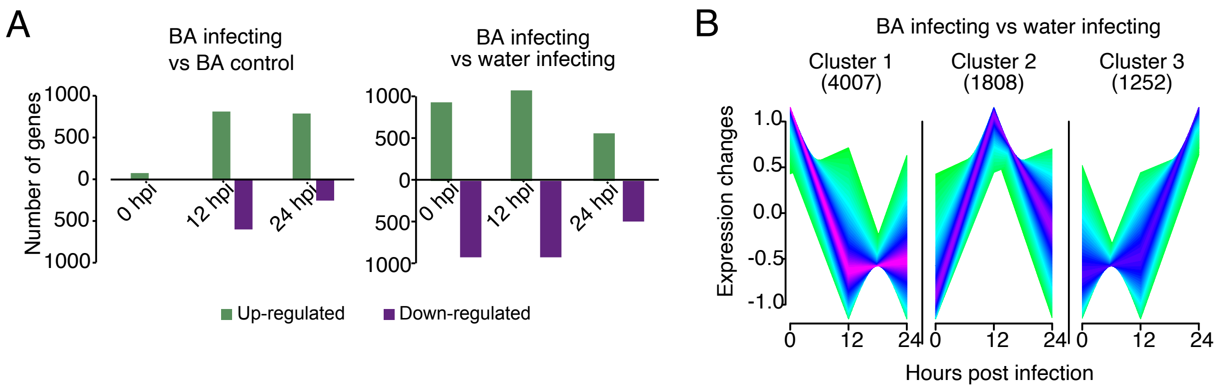


C

| Cluster 1 (BA infect vs water infect) | | | Cluster 2 (BA infecting vs water infecting) | | | Cluster3 (BA infect vs water infect) | | |
|---|---|---|---|---|---|---|---|---|
| GO term | | p-value | GO term | | p-value | GO term | | p-value |
| GO:0006468 | protein phosphorylation | 1.20E-09 | GO:0006260 | DNA replication | 3.30E-20 | GO:0055114 | oxidation-reduction process | 0.003 |
| GO:0016567 | protein ubiquitination | 0.00024 | GO:0000105 | histidine biosynthetic process | 3.60E-05 | GO:0006783 | heme biosynthetic process | 0.0041 |
| GO:0005992 | trehalose biosynthesis | 0.002 | GO:0006302 | double-strand break repair | 6.80E-05 | GO:0015074 | DNA integration | 0.006 |
| GO:1903506 | regulation transcription | 0.00312 | GO:0009451 | RNA modification | 0.00012 | GO:0006508 | proteolysis | 0.0085 |
| GO:0015770 | sucrose transport | 0.00873 | GO:0016072 | rRNA metabolic process | 0.00031 | GO:0006450 | regulation of translational fidelity | 0.0443 |
| GO:0006680 | glucosylceramide catabolism | 0.01807 | GO:0022613 | ribonucleoprotein biogenesis | 0.00038 | GO:0009229 | thiamine diphosphate biosynthesis | 0.0443 |
| GO:0048268 | clathrin coat assembly | 0.03064 | GO:0006457 | protein folding | 0.00079 | GO:0032780 | negative regulation of ATPase activity | 0.0443 |
| GO:0016573 | histone acetylation | 0.03494 | GO:0009089 | lysine biosynthetic process | 0.00082 | | | |
| GO:0023052 | signaling | 0.03505 | GO:0006310 | DNA recombination | 0.00097 | | | |
| GO:0000288 | mRNA catabolism | 0.041 | GO:0006418 | tRNA aminoacylation | 0.00107 | | | |
| GO:0006665 | sphingolipid metabolism | 0.04925 | GO:0006207 | de novo' pyrimidine biosynthesis | 0.00333 | | | |
| | | | GO:0032259 | methylation | 0.00472 | | | |

D

| BA infecting vs BA control up-regulated | | | | | | | | |
|---|---|---|---|---|---|---|---|---|
| 0 hpi | | p-value | 12 hpi | | p-value | 24 hpi | | p-value |
| GO:0009690 | cytokinin metabolic process | 3.30E-08 | GO:000969 | cytokinin metabolic process | 1.90E-06 | GO:0009690 | cytokinin metabolic process | 5.00E-06 |
| GO:0006855 | drug transmembrane transport | 2.20E-07 | GO:0006096 | glycolytic process | 4.00E-05 | GO:0042744 | hydrogen peroxide catabolic process | 5.50E-06 |
| GO:0000160 | phosphorelay signal transduction system | 1.50E-06 | GO:0006108 | malate metabolic process | 0.00012 | GO:0071555 | cell wall organization | 7.30E-05 |
| GO:0009664 | plant-type cell wall organization | 0.00012 | GO:0000160 | phosphorelay signal transduction system | 0.00015 | GO:0055114 | oxidation-reduction process | 0.00012 |
| GO:0042744 | hydrogen peroxide catabolic process | 0.00013 | GO:0055114 | oxidation-reduction process | 0.00021 | GO:0006979 | response to oxidative stress | 0.00018 |
| GO:0006979 | response to oxidative stress | 0.00029 | GO:0006006 | glucose metabolic process | 0.00022 | GO:0045492 | xylan biosynthetic process | 0.00024 |
| GO:0015770 | sucrose transport | 0.00628 | GO:0051187 | cofactor catabolic process | 0.00044 | GO:0008272 | sulfate transport | 0.00219 |
| GO:0005986 | sucrose biosynthetic process | 0.03312 | GO:0009607 | response to biotic stimulus | 0.00074 | GO:0006869 | lipid transport | 0.0023 |
| GO:0006006 | glucose metabolic process | 0.0025 | GO:0006855 | drug transmembrane transport | 0.00099 | GO:0042546 | cell wall biogenesis | 0.00484 |
| GO:0009688 | abscisic acid biosynthetic process | 0.03312 | GO:0042737 | drug catabolic process | 0.00099 | GO:0010105 | negative regulation of ethylene-activate... | 0.04928 |

| BA infecting vs BA ccontrol down-regulated | | | | | | | | |
|---|---|---|---|---|---|---|---|---|
| 0 hpi | | p-value | 12 hpi | | p-value | 24 hpi | | p-value |
| GO:0006857 | oligopeptide transport | 0.00058 | GO:0046274 | lignin catabolic process | 1.00E-05 | GO:0007018 | microtubule-based movement | 4.10E-07 |
| GO:0070588 | calcium ion transmembrane transport | 0.00214 | GO:2001141 | regulation of RNA biosynthetic process | 1.50E-05 | GO:0055085 | transmembrane transport | 0.00099 |
| GO:0048544 | recognition of pollen | 0.00308 | GO:0048544 | recognition of pollen | 0.00049 | GO:0006857 | oligopeptide transport | 0.00469 |
| GO:0009607 | response to biotic stimulus | 0.00365 | GO:0055114 | oxidation-reduction process | 0.00061 | GO:0030001 | metal ion transport | 0.0218 |
| GO:0055114 | oxidation-reduction process | 0.00494 | GO:0070588 | calcium ion transmembrane transport | 0.00203 | GO:0000165 | MAPK cascade | 0.0296 |
| GO:0006952 | defense response | 0.00659 | GO:0009607 | response to biotic stimulus | 0.00337 | GO:0005992 | trehalose biosynthetic process | 0.03229 |
| GO:0008283 | cell proliferation | 0.00896 | GO:0006950 | response to stress | 0.00409 | GO:0048544 | recognition of pollen | 0.0331 |
| GO:0042744 | hydrogen peroxide catabolism | 0.00949 | GO:0006020 | inositol metabolic process | 0.00529 | GO:0055114 | oxidation-reduction process | 0.04856 |
| GO:0000154 | rRNA modification | 0.01317 | GO:0000165 | MAPK cascade | 0.01128 | | | |
| GO:0006809 | nitric oxide biosynthetic process | 0.03092 | GO:0006809 | nitric oxide biosynthetic process | 0.03035 | | | |

E

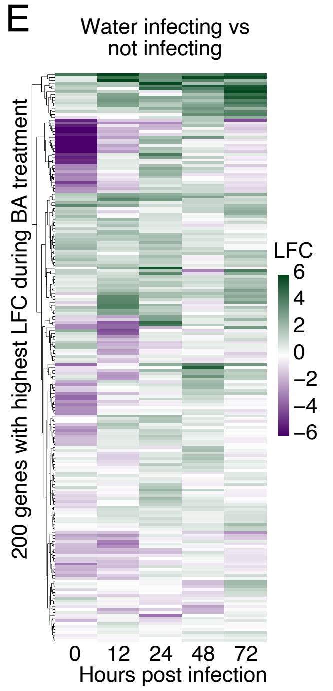


F

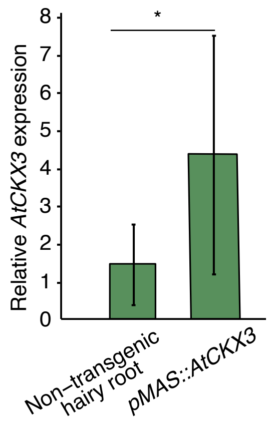


**Figure S4: Exogenous cytokinin treatment induces gene expression changes in *Phtheirospermum***

**A)** Number of differentially expressed genes over three time points in the BA infecting versus BA control and BA infecting versus water infecting RNA-seq libraries in *Phtheirospermum*. **B)** Clustering of differentially expressed (DE) genes in BA infecting versus water infecting RNA-seq datasets of *Phtheirospermum* infecting *Arabidopsis* over three time points based on their co-expression patterns. The number in parenthesis is the number of genes in each cluster. **C)** Gene ontology analysis for the DE genes assigned to each co-expression cluster for the BA infecting versus water infecting RNA-seq datasets (n = 3 libraries, Wald test with Benjamini-Hochberg correction, p<0.05). **D)** Gene ontology analysis for the up or down regulated genes for the BA infecting versus BA control RNA-seq datasets (n = 3 libraries, Wald test with Benjamini-Hochberg correction, p<0.05). **E)** Heatmap of the 200 genes with the highest log2 fold change (LFC) after BA treatment shown over five time points in water infecting versus not infecting *Arabidopsis* roots RNAseq libraries. **F)** qRT-PCR gene expression quantification of *AtCKX3* in *Phtheirospermum* transgenic or non-transgenic hairy roots. (n = 5-6 replicates, Student's t-test, * p<0.05).
